# NEW INSIGHTS INTO THE HEPATIC IRON PHENOTYPE OF BMP6 KNOCKOUT MICE

**DOI:** 10.1101/2023.09.28.559941

**Authors:** Céline Besson, Alexandra Willemetz, Chloé Latour, Lorenne Robert, Helene Coppin, Marie-Paule Roth, François Canonne-Hergaux

## Abstract

**Objective:** *Bmp6* knockout (KO) mice progressively accumulate a significant amount of iron in their liver as they age due to a defect in hepcidin (Hamp) expression and an upregulation of the iron exporter ferroportin (Fpn). In this study, we conducted a comprehensive investigation of the hepatic iron overload phenotype, with specific emphasis on the cellular and subcellular localization of Fpn in *Bmp6* KO mice.

**Materials and Methods:** Livers obtained from *Bmp6* knockout (KO) mice at different developmental stages were utilized for the quantification of iron content, investigation of iron distribution, histological analysis, histoimmunofluorescence assays performed on paraffin-embedded sections, confocal microscopy examinations, subcellular membrane fractionation, and western blot analysis.

**Results:** In *Bmp6* KO livers, iron overload increased with age and was not homogeneous, with certain hepatic lobes and specific areas in liver sections showing more pronounced iron accumulation. In young mice, iron accumulated mostly in the centrilobular zone where low Fpn expression was observed. Fpn was strongly detected in periportal Kupffer cells and at the apical membrane of periportal hepatocytes lining the sinusoidal capillaries. The zonal distribution of iron tended to disappear with age in strongly iron-overloaded areas, with the appearance of large cellular aggregates strongly positive for Fpn, iron, and ceroid/lipofuscin. At the subcellular level, hepatic Fpn seemed to concentrate in specific cell surface compartments and was enriched in a lipid raft fraction.

**Conclusions:** Unregulated expression of Fpn on the cell surface of periportal macrophages and hepatocytes results in centrilobular iron overload within hepatocytes. In areas of pronounced iron overload, Fpn expression is present in lipogranulomas, identified as aggregations of macrophages accumulating hemosiderin and ceroid/lipofuscin pigments. These lesions likely form due to the phagocytosis of sideronecrotic/ferroptotic hepatocytes by macrophages. In contrast to the duodenal form of Fpn, both splenic and hepatic Fpn demonstrated robust enrichment within lipid rafts. The observed variations in the subcellular localization of Fpn could play a significant role in influencing the transporter’s iron transport activity and/or its regulation by hepcidin.

## INTRODUCTION

*Bmp6* knockout (KO) mice progressively accumulate a significant amount of iron in their liver as they age due to a defect in hepcidin (Hamp) expression and a substantial upregulation of the iron exporter ferroportin (Fpn, also referred to as Slc40a1, solute carrier family 40 member 1) ^1,2^. Iron deposits in the *Bmp6* KO liver have been shown to be more pronounced in the centrilobular areas ^2^. However, the liver, being a complex organ, possesses a distinctive anatomy characterized by distinct lobes that exhibit morphological and functional variations ^3^. In fact, limited information is available concerning the precise distribution of iron in histologic sections throughout the hepatic lobes of these mice. As a result, we undertook a meticulous examination of iron overload in each hepatic lobe of *Bmp6* KO mice and tracked its changes with advancing age. Furthermore, we investigated the expression and localization of Fpn in the liver, in conjunction with the distribution of iron deposits.

In *Bmp6* KO mice, Fpn was described to be overexpressed in enterocytes of the duodenum, in splenic macrophages and cells with in the liver ^2^. A similar phenotype was observed in *Hamp* KO mice ^4^. However, the hepatic cellular localization of Fpn was only partially characterized in both *Hamp* and *Bmp6* KO mice, using immunohistochemistry techniques ^2,4,5^. Interestingly, in *Hamp* KO mice, Fpn was found to be more expressed in periportal areas ^5^. Such zonal tissue distribution has yet to be explored in *Bmp6* KO. Previously, in both *Bmp6* and *Hamp* KO liver, Fpn was detected in Kupffer cells and in the perisinusoidal space (aka space of Disse) lining the apical membranes of hepatocytes ^4,5,5^. While hepatocytes were also demonstrated to express Fpn ^5^, the exact subcellular localization of the iron exporter within these cells remains unknown. Furthermore, beyond parenchymal hepatocytes, other cell types such as sinusoidal endothelial cells (SEC) and hepatic stellate cells (HSC) coexist within the space of Disse. Notably, mRNA studies have indicated substantial Fpn expression in HSC within the rat liver ^6^, suggesting a plausible presence of the iron transporter within these cells.

Consequently, we conducted a meticulous analysis of the expression and the localization of Fpn at the tissue, cellular, and subcellular levels in *Bmp6* KO mice. This investigation involved immunofluorescence studies, colocalization labeling, and subcellular fractionation techniques.

## MATERIALS AND METHODS

### Animals

Animal experiments were carried out following approved conditions and protocols by the local ethics committee (UMS006 CEEA-122 Toulouse) and in accordance with the European Union directive 2010/63/EU. The generation of *Bmp6*^tm1Rob^ mice (*Bmp6^−/−^*: *Bmp6* KO) on an outbred CD1 background was previously documented ^7^. Mice were genotyped using high-resolution amplicon melting via the LightCycler 480 System (Roche Diagnostics). Wildtype and *Bmp6* KO mice (only males) were euthanized at various ages, as specified, and tissues were collected for the different analyses.

### Antibodies sources

To detect Fpn, experiments were conducted using either with an antibody purchased from Alphadiagnostic (MTP11-A; immunofluorescence) or with a homemade produced and affinity-purified polyclonal rabbit antisera against Fpn (immunofluorescence and western blotting) ^8^. Polyclonal rabbit antisera against Dmt1 (Slc11a2 solute carrier family 11 member 2, aka as Nramp2) were generated and affinity purified as described previously ^9^. The specificity of homemade Fpn and Dmt1 sera was validated by immunoblotting, immunofluorescence and histochemistry ^2,8,9^. The anti-human caveolin-1 (Cav1) antibody was purchased from Tebu-bio (N-20). The rabbit anti-heme-oxygenase-1 (Hmox1) polyclonal antibody (SPA-896) was purchased from Stressgene Biotechnology. The mouse monoclonal anti-vinculin antibody (VINC) was purchased from Sigma–Aldrich (CLONE hVIN-1). The rat monoclonal anti-mouse dipeptidyl peptidase-4 (Dpp4 also known as CD26) and the rat monoclonal anti-mouse F4/80 antibodies were purchased from Abcam (ab42899) and AbD serotec (CL:A3-1), respectively. The goat polyclonal anti-Desmin antibody (Y-20) was purchased from Santa Cruz (sc-7559). Secondary peroxidase antibodies against mouse, rat and rabbit were obtained from DAKO. The secondary antibodies used for immunofluorescence, including Alexa Fluor® 488 goat anti-rabbit Ig (GAR-alexa488), Alexa Fluor® 568 Rat anti-mouse Ig (RAM-alexa568), and Alexa Fluor® 563 goat anti-rat Ig (GARa-alexa563), were obtained from Molecular Probes.

### Tissue iron staining and quantitative iron measurement

Left lateral lobes (LLL), right lateral lobes (RLL), medium lobes (ML) and caudate lobes (CL) of the liver from three mice per genotype were fixed in 4% buffered formalin and subsequently embedded in paraffin. Following deparaffinization, tissue sections were subjected to staining using Perls Prussian blue stain to visualize nonheme iron, followed by counterstaining with nuclear fast red. The slides were observed under a light microscope and images were captured or scanned using a Pannoramic 250 Flash II (3DHISTECH). Subsequent analysis was performed utilizing a Pannoramic Viewer software. Due to the size of certain lobes, quantitative assessments of nonheme iron in hepatic lobes were conducted by pooling each specific lobes from three mice per genotype (pooled lobes: 3xLLL, 3xRLL, 3xML or 3xCL). Iron measurements of the pooled lobes were performed following the methodology recommended by Torrance and Bothwell ^10^. Briefly, 100 mg of tissues were homogenized in 150 µl water using a FastPrep-24 Instrument (MP Biomedicals) for 60 seconds at 6 m/second. The homogenates were then mixed with a solution containing HCl 30%/trichloroacetic acid 10% to achieve a final volume of 1.5 ml. These samples were allowed to incubate overnight at 65°C. The iron content was subsequently complexed with the bathophenantroline sulfonate chromogen, and the absorbance was measured at a wavelength of 535 nm.

### Histology and immunohistofluorescence analysis

Following fixation and embedding in paraffin, each lobe (LLL, RLL, ML, and CL) was prepared for tissue sectioning. Following blocking with BSA 1% and 10% heat inactivated goat serum for 30 min at room temperature, deparaffinized liver sections were incubated with primary antibodies for 1hr. After three washes with PBS containing 0.5% BSA, the sections were incubated with Alexa conjugated secondary antibodies for an additional one hour at room temperature. Once mounted, sections were visualized using an epifluorescence microscope LEICA DM-IRM, a Zeiss confocal fluorescent microscope or a Pannoramic 250 Flash II (3DHISTECH) scanner. Image acquisition was performed using either ARCHIMED-PRO (Microvision Instruments), Zeiss LSM Image Browser softwares or the Pannoramic Viewer software.

### Preparation of crude membrane and cytosolic protein extracts

Cytosolic and membrane fractions were obtained following a previously established and validated protocol ^8^. In brief, tissue samples were homogenized using an Ultraturax in lysis buffer (10 mL/g of tissue in 0.25 mol/L sucrose/0.03 mol/L histidine, pH 7.2) which was supplemented with protease inhibitors (PI), and phenylmethylsulfonyl fluoride (PMSF). The resulting lysates were centrifuged at 6000 g for 15 min to remove nuclei and unbroken cells. The resulting supernatants, termed post-nuclear supernatants (PNS) were subsequently subjected to ultracentrifugation (Beckman) at 80,000 g for 30 minutes to separate the crude membrane fractions from the cytosolic proteins. Supernatants corresponding to cytosolic extracts were collected and the membrane pellets were resuspended in a lysis buffer (0.25 mol/L sucrose/0.03 mol/L histidine, pH 7.2), supplemented with PI’s, and PMSF. The protein concentrations of both the membrane and cytosolic fractions were determined via the Bradford assay (Bio-RAD) and the fractions were stored at –80°C until utilization.

### Isolation of lipid rafts/detergent resistant membranes

Lipid rafts from mouse tissues were isolated as detergent (Triton X-100) resistant membranes [DRM, ^11^]. Grounded frozen tissues (liver, spleen and duodenum) were used to prepare crude membrane extracts as described before ^8^. Membrane pellets were resuspended in lysis buffer (1ml/pellet from 100mg tissues; 150 mM NaCl, 25 mM MES, 5 mM EDTA, pH 6.5, 1% Triton X-100) supplemented with protease inhibitor cocktail (PI’s; Sigma Aldrich) and phenylmethanesulfonyl fluoride (PMSF; Sigma Aldrich). Samples were then homogenized using a mini plastic potter and tissue lysates were incubated at 4°C for 60 min with gentle agitation. Samples were then homogenized by 20 passages through a 25-gauge needle (5/8-inch) and adjusted to a final concentration of 40% (w/v) Iodixanol (OptiPrep®, sigma Aldrich). The mixture was then layered under a 20–40% discontinuous iodixanol gradient and centrifuged at 260,000 g for 15h at 4°C using an SW 41 Ti Rotor (Beckman Coulter). After spinning, twelve fractions of 1 ml were collected from the top to the bottom of the gradient tube and the protein concentration of each fraction was determined with a Bradford assay before immunoblotting.

### Immunoblotting

Proteins samples (crude membrane extracts and DRM) were prepared in Laemmli buffer (50mM Tris, 55mM SDS, 1M Glycerol, 570mM β-mercaptoethanol and bromophenol blue) and incubated for 30 min at RT. Samples were separated in a 10% SDS-PAGE and transferred onto PVDF membrane. To control protein loading and transfer, membranes were stained with 0,1% Red Ponceau S. Membranes were then distained in 0,1% Tween-PBS (T-PBS) and blocked O/N with 5% skim milk in T-PBS at 4°C or for 2h at RT. Each membrane was incubated with primary antibodies diluted in blocking solution for 2h at RT or O/N at 4°C. After washing in T-PBS, blots were incubated for 1h at RT with the secondary antibodies conjugated with horseradish peroxidase (HRP) diluted in blocking solution. Membranes were washed in T-PBS and followed by development with either Novex ECL (Invitrogen) or Immobilon Western (Millipore) Chemiluminescent HRP substrate reagents.

### RNA studies

Total RNA from the liver was extracted using Isol-RN lysis reagent (5 PRIME). Complementary DNA was synthesized using MMLV-RT (Promega). The sequence of the primers for the *Hamp*, *Fpn*, and the reference gene *Hprt* are listed in Supporting Table S1. Quantitative polymerase chain reactions (PCRs) were prepared with LightCycler 480 DNA SYBR Green I Master reaction mix and run on a LightCycler 480 System (both from Roche Diagnostics). Cycle threshold difference (DCt) values were obtained by subtracting the reference gene Ct value to the target gene Ct. GraphPad Prism (version 6) was used for statistical analyses (GraphPad Software, La Jolla California USA) and DCt values in WT and *Bmp6* KO mice were compared by Student’s t-Tests.

## RESULTS

### Ferroportin and *Hamp1* expression in *Bmp6* KO liver

We assessed the hepatic expression of Fpn and Hamp mRNA through RT-qPCR in both WT and *Bmp6* KO mice (Fig.1A). Consistent with prior findings ^2^, the Hamp mRNA levels exhibited a substantial decrease in *Bmp6* KO mice compared to their WT counterparts. On the other hand, *Fpn* mRNA expression did not change significantly. In contrast to these observations, the protein level of Fpn was notably elevated in *Bmp6* KO mice, as demonstrated by western blot analysis (Fig.1B) and immunohistofluorescence of Fpn within liver sections (Fig.1C).

**Fig.1:**
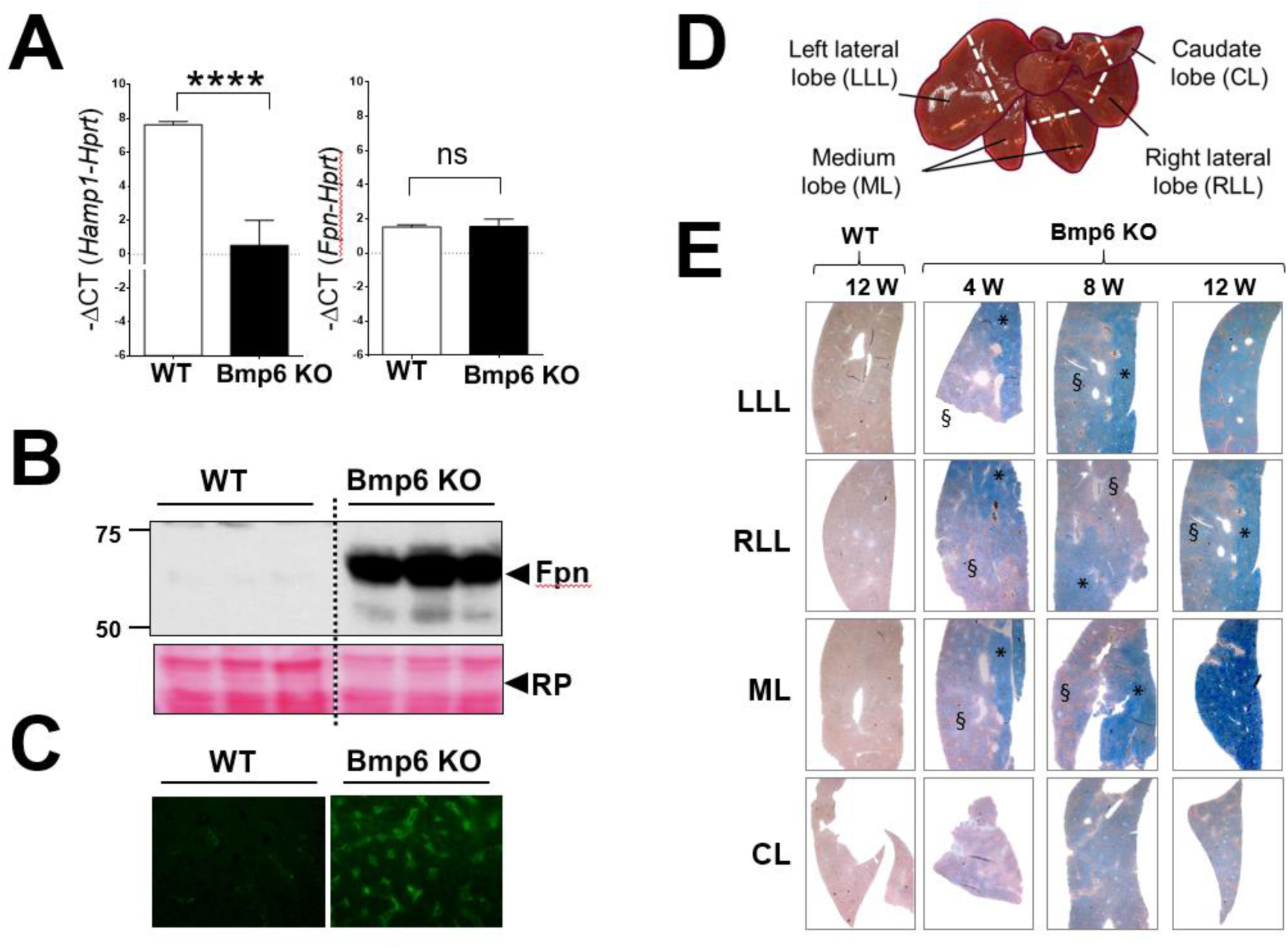
Expression of ferroportin (Fpn) and hepcidin 1 (Hamp1), and iron overload in Bmp6 KO liver. (A) Messenger RNA expression levels of hepcidin (Hamp) and ferroportin (Fpn) in total liver samples from wildtype (WT) or *Bmp6* KO mice (n=6). The relative gene expression is presented as –ΔCT (CT gene of interest - CT Hprt). Statistical significance: P<0.0001 (****), ns: not significant. (B) Total membrane proteins were extracted from WT or *Bmp6* KO livers (three mice per genotype) and subjected to Western blot analysis. Protein visualization was assessed both quantitatively and qualitatively using Ponceau S staining (PS). Molecular weight markers and their corresponding sizes in kilodaltons (kDa) are indicated. (C) Immunofluorescence validation of the substantial upregulation of Fpn protein in liver sections. (D) Illustration of the different parts of the distinct lobes [left lateral lobe (LLL), right lateral lobe (RLL), medium lobe (ML), and caudate lobe (CL)] used for tissue sectioning and other analyses. (E) Representative images (of three mice per genotype) illustrating iron distribution in liver lobes [ (LLL), (RLL), (ML), and (CL)] from wildtype (WT) and *Bmp6* KO mice at different ages. Perls staining was performed for iron localization. In *Bmp6* KO mice (B), iron distribution was not uniform among lobes or within the same lobe, with certain zones displaying lower (§) and higher iron accumulation (*).

### Hepatic iron localisation in *Bmp6* KO liver

Subsequently, we proceeded to track the iron accumulation pattern across distinct liver lobes in both wildtype mice (WT) and *Bmp6* KO mice (Fig.1D). The left lateral lobe (LLL), right lateral lobe (RLL), medium lobe (ML), and caudate lobe (CL) were subjected to Perls staining and analyzed via microscopy. Among 12-week-old WT mice (CD1), there were no prominent indicators of iron accumulation within their liver lobes. A meticulous examination revealed slight iron loading in specific periportal regions (see Fig.2A). Importantly, no significant or marked differences in iron loading were observable in WT mice from 4 weeks to 12 weeks (not shown). In contrast to WT mice, *Bmp6* KO mice displayed a significant iron overload even at the age of 4 weeks (Fig.1D). In these mice, iron overload increased as they aged in each of the liver lobes

**Fig.2:**
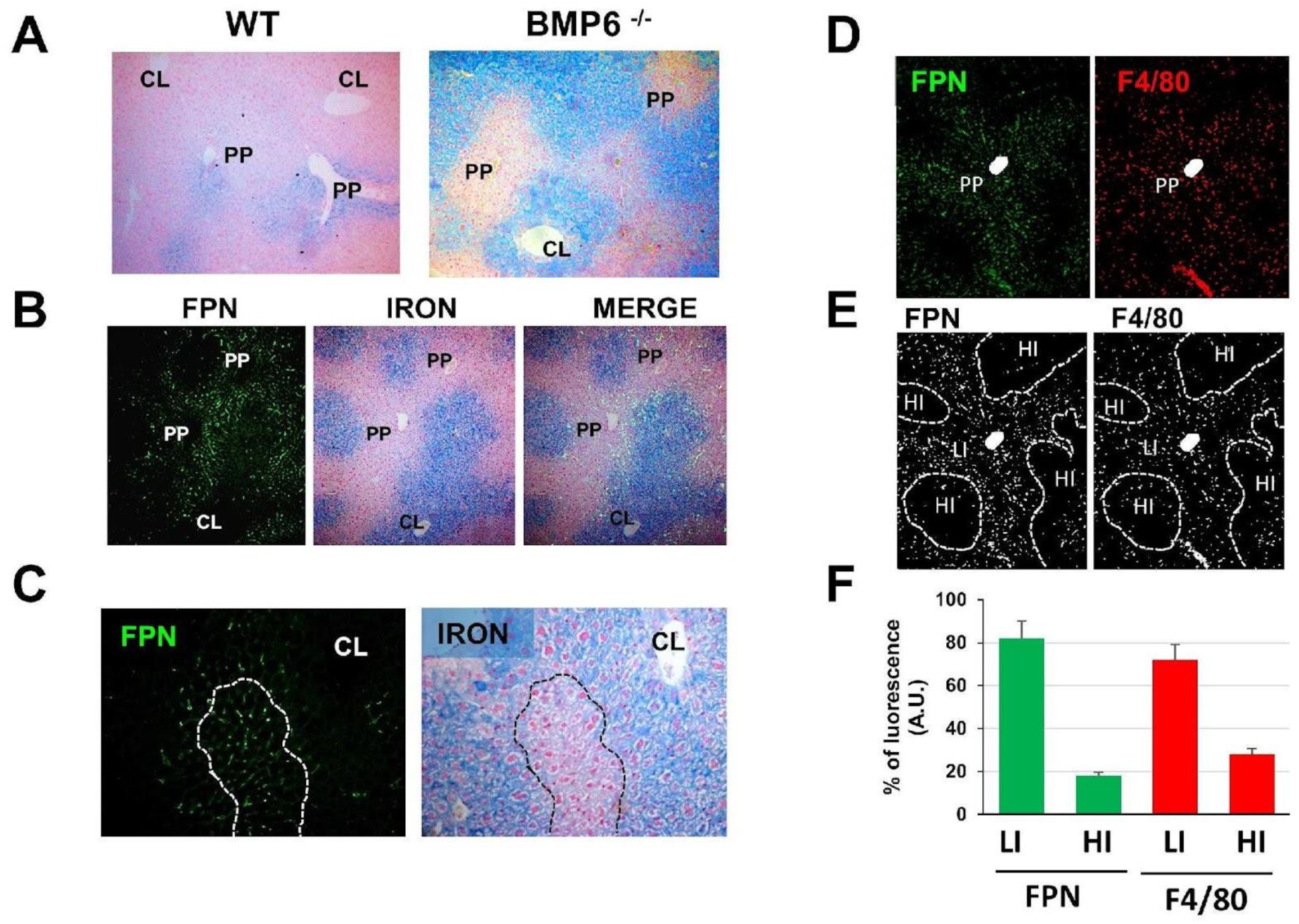
Iron, ferroportin (Fpn), and macrophage distribution in *Bmp6* KO liver from young mice. **(A)** Perls staining of hepatic left lateral lobe (LLL) from 8-week-old wildtype (WT) and *Bmp6* knockout (KO) mice. Images for *Bmp6* KO mice in (A) depict zones with mild iron accumulation, marked as (§). In contrast to wildtype mice showing some iron in the periportal zone (PP), *Bmp6* KO mice exhibited iron accumulation in the centrilobular zone (CL). **(B and C)** Overlay of Perls staining images with Fpn immunofluorescence visualization, indicating the concentration of Fpn staining in periportal low iron zones. **(D to F)** Immunofluorescence analysis of Fpn and F4/80 (macrophage marker) staining. **(F)** Histogram representation of Fpn and F4/80 fluorescence (% of all fluorescence dots) from the images in panel E. Fpn and F4/80 were enriched in the same zones corresponding to periportal low iron zones. LI: Low iron zones; HI: High iron zones

However, the distribution of iron deposits was not homogenous between hepatic lobes, with the ML exhibiting the highest iron accumulation over time. Such observation was corroborated by the quantitative measurements of iron in each lobe of *Bmp6* KO mice (Fig.S1). Interestingly, this variation in iron accumulation among lobes was not accompanied by changes in Hamp and Fpn mRNA expression (not shown). Importantly, within each lobe, except for the caudate lobe (CL), certain specific areas [Fig.1D (*)] exhibited more pronounced iron accumulation compared to other regions [Fig.1D (§)].

### Histological analysis of iron-rich hepatic regions in young and aged *Bmp6* KO mice

Concurrently, we examined the pattern of Fpn expression alongside the distribution of cellular iron overload in the livers of young *Bmp6* KO mice (Fig.2). As discussed earlier, under high magnification, a subtle iron accumulation was discernible in specific hepatocytes within the periportal area of WT (CD1) mice (Fig.2A, PP). In contrast, young *Bmp6* KO mice displayed predominant iron accumulation in hepatocytes localized within the centrilobular zones (Fig.2A, B, & C; CL), where a lower Fpn expression was observed (Fig.2B and C). Specifically, the expression of Fpn was largely confined to cells localized in the periportal area (Fig.2C & D). Histograms depicting the fluorescence of Fpn and F4/80 (a macrophage marker) (% of total fluorescence dots, Fig.2F) from the images presented in panels D and E demonstrated that both Fpn and F4/80 exhibited enrichment in the same regions corresponding to periportal zones associated with lower iron levels.

In areas of pronounced iron overload [Fig.1D and Fig.S2 (*)] which became more prominent with advancing age, particularly within the medium lobe (ML), the characteristic zonal distribution of iron disappeared with the appearance of large aggregates strongly enriched in iron (Fig.3A, arrows; Fig.S2 A and A’, arrowheads). Such aggregates were shown to be positive for both Fpn (Fig.3B and C; Fig.S2 B and B’; green fluorescence) and Soudan black (Fig.3B and C; Fig.S2 C and C’, brown coloration). Furthermore, these aggregates exhibited autofluorescence (Fig.3D, in red) and were also positive for F4/80 (Fig3E). In comparison to other liver lobes, the CL seemed to exhibit a greater resistant to strong iron accumulation (Fig.1 & S1) and did not display cellular aggregates (Fig.S2D).

**Fig.3:**
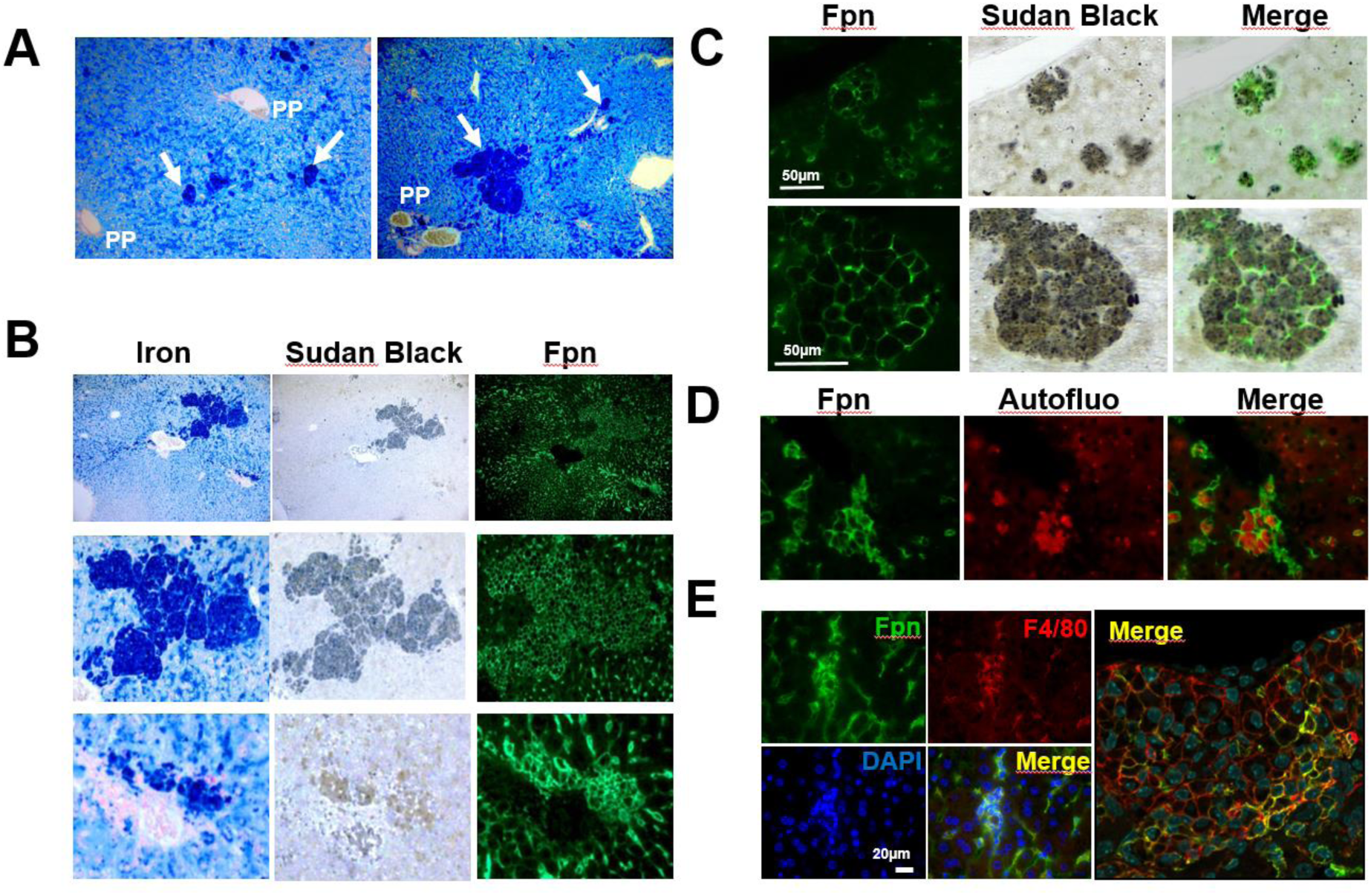
Histology and immunofluorescence analysis of high iron-containing zones identified in *Bmp6* KO liver. **(A)** Perls staining of a medium lobe (ML) from Bmp6 KO mice at 12 weeks of age. The zonal iron distribution is disrupted in regions with significant iron overload, displaying the emergence of prominent iron aggregates (arrow). **(B)** Sudan black (ceroid/lipofuscin) and Fpn immunofluorescence staining in areas of intense iron overload (Perls staining) present in the left lateral lobe (LLL) from a 12-week-old KO *Bmp6* Mice. **(C)** Ferroportin staining strongly correlates with ceroid/lipofuscin accumulation. **(D)** Ferroportin staining is associated with autofluorescence within the aggregates. **(E)** Merged colabeling of Fpn and F4/80 highlights the presence of numerous macrophages within these cellular aggregates.

Collectively, these observations suggest that within liver regions characterized by substantial iron accumulation, the emergence of iron-positive cellular aggregates primarily consists of macrophages. These macrophages exhibit strong expression of Fpn and contain ceroid/lipofuscin (identified by Soudan black staining, as further discussed).

### Cellular and subcellular localisation of ferroportin in *Bmp6* KO liver

To precisely ascertain the subcellular localization of Fpn, we conducted a detailed confocal analysis within liver sections from 8-week-old *Bmp6* KO mice, involving co-immunofluorescence staining of the iron exporter Fpn alongside markers such as F4/80 (macrophage marker), CD26 (also known as dipeptidyl peptidase-4, Dpp-4, expressed both in the apical membrane of hepatocytes and in hepatocyte plasma membrane forming the biliary canaliculi), and desmin (a stellate cell marker) (Fig. 4). Fpn was strongly detected in periportal Kupffer cells (Fig.4A and A’, arrowhead) as well as at the apical membrane of hepatocytes lining the sinusoidal capillaries (arrows in Fig.4A, A’ and 4B, B’). However, Fpn was not detected in the Dpp4 positive membrane of the biliary canaliculi (arrowhead in Fig.4B, B’). The absence of colocalization between Fpn and desmin (Fig.3C, C’) suggested that stellate cells did not express Fpn. In addition, there was no detection of Fpn in endothelial CD34-positive cells (not shown).

**Fig.4:**
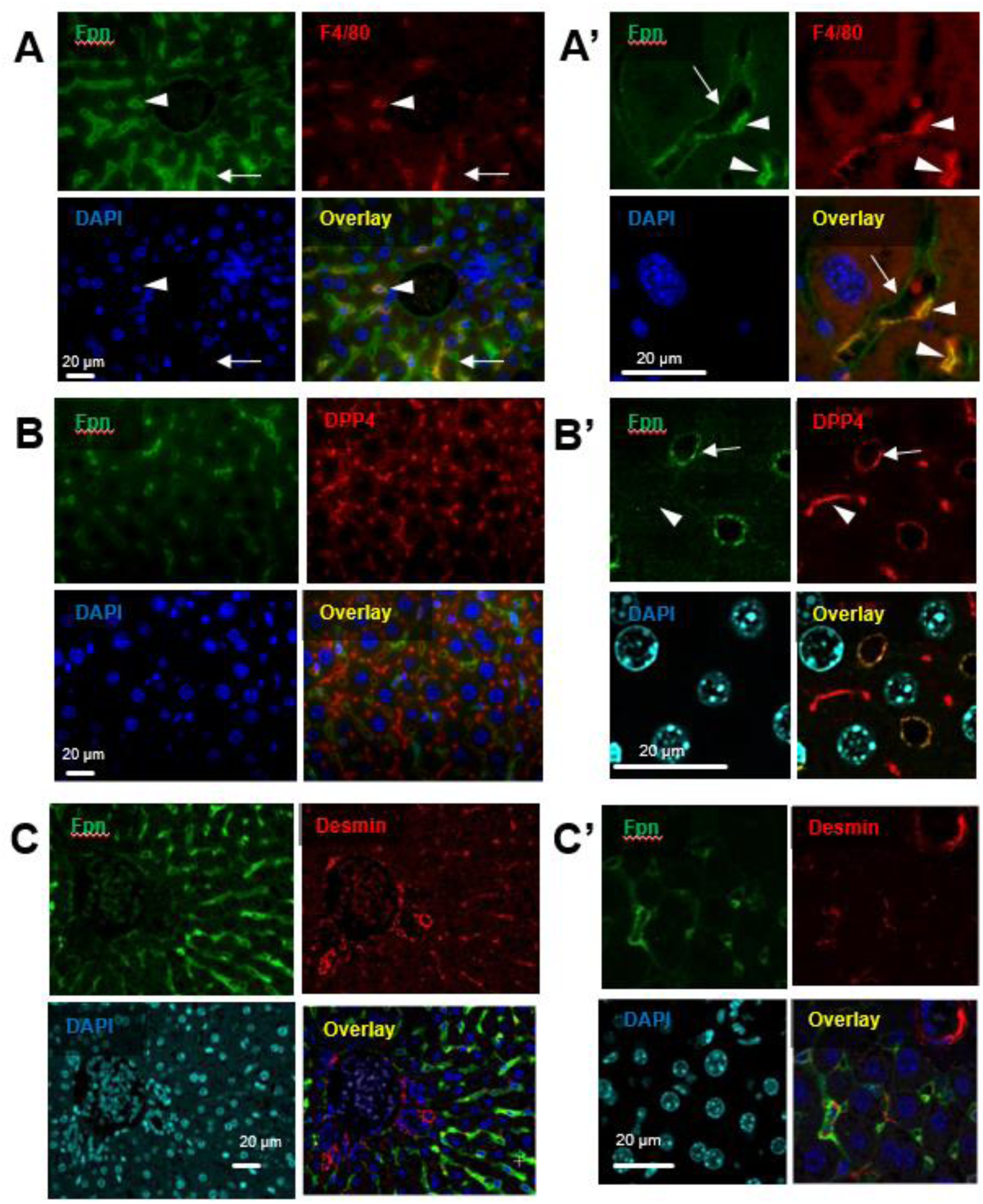
Confocal analysis of co-immunofluorescence staining of ferroportin (Fpn) with F4/80 (A & A’), Dipeptidyl peptidase-4 (DPP-4, B & B’) or Desmin (C & C’) in liver sections of 8 weeks-old *Bmp6* KO mice. Orange/yellow color in the overlays (A, A’ and B, B’) indicates the colocalization of Fpn with F4/80 and DPP-4. No such colocalization of Fpn was observed with Desmine. Nuclear DNA labeling was achieved using DAPI.

Interestingly, the hepatic Fpn detected in *Bmp6* KO mice appeared to concentrate within specific membrane compartments at the cell surface of both hepatocytes (arrows) and Kupffer cells (arrowheads in Fig.5A). Notably, Dpp4, which colocalised with Fpn in apical membrane of hepatocytes (Fig.5B), is known to be enriched in lipid rafts ^12,13^. Previous observations in cultured macrophages have clearly demonstrated that Fpn is mostly present in lipid rafts ^11^. Therefore, we decided to investigate the subcellular localisation of Fpn within lipid rafts in *Bmp6* KO tissues, namely the liver, spleen, and duodenum, using the detergent-rich membrane (DRM) analysis (see M&M and Fig.S4). As previously described ^2^, Fpn was also strongly increased in spleen and duodenum of *Bmp6* KO mice (Fig.S3). Co-staining of F4/80 and Fpn in the spleen indicated that Fpn was in macrophages of the red pulp (Fig.5C). In the duodenum, Fpn was strongly expressed at the basolateral membrane of enterocytes, in sharp contrast to the brush border localization of the iron transporter Dmt1 (Fig.5C). Using iodixanol density gradient centrifugation and western blot analysis (Fig.S4), we compared the subcellular fractionation of hepatic Fpn with those obtained from splenic and duodenal Fpn of 8 week-old *Bmp6* KO mice (Fig.5D and E). Similar volumes (Fig.5D) or similar amounts of protein (Fig.5E) from pooled fractions F1, F2, F3 and F4 (see M&M and Fig.S4) were analyzed by western blotting. Interestingly, hepatic Fpn as well as splenic Fpn were strongly enriched in the fraction containing the lipid raft marker caveolin 1 (Cav1). As a control and in contrast to Fpn, Hmox1 (Heme oxygenase 1) was mainly detected in Non Detergent resistant membrane (F4). On the other hand, duodenal Fpn as well as Dmt1 were mainly detected in non-raft (Cav1 negative) containing fractions F3 and F4, respectively.

**Fig.5:**
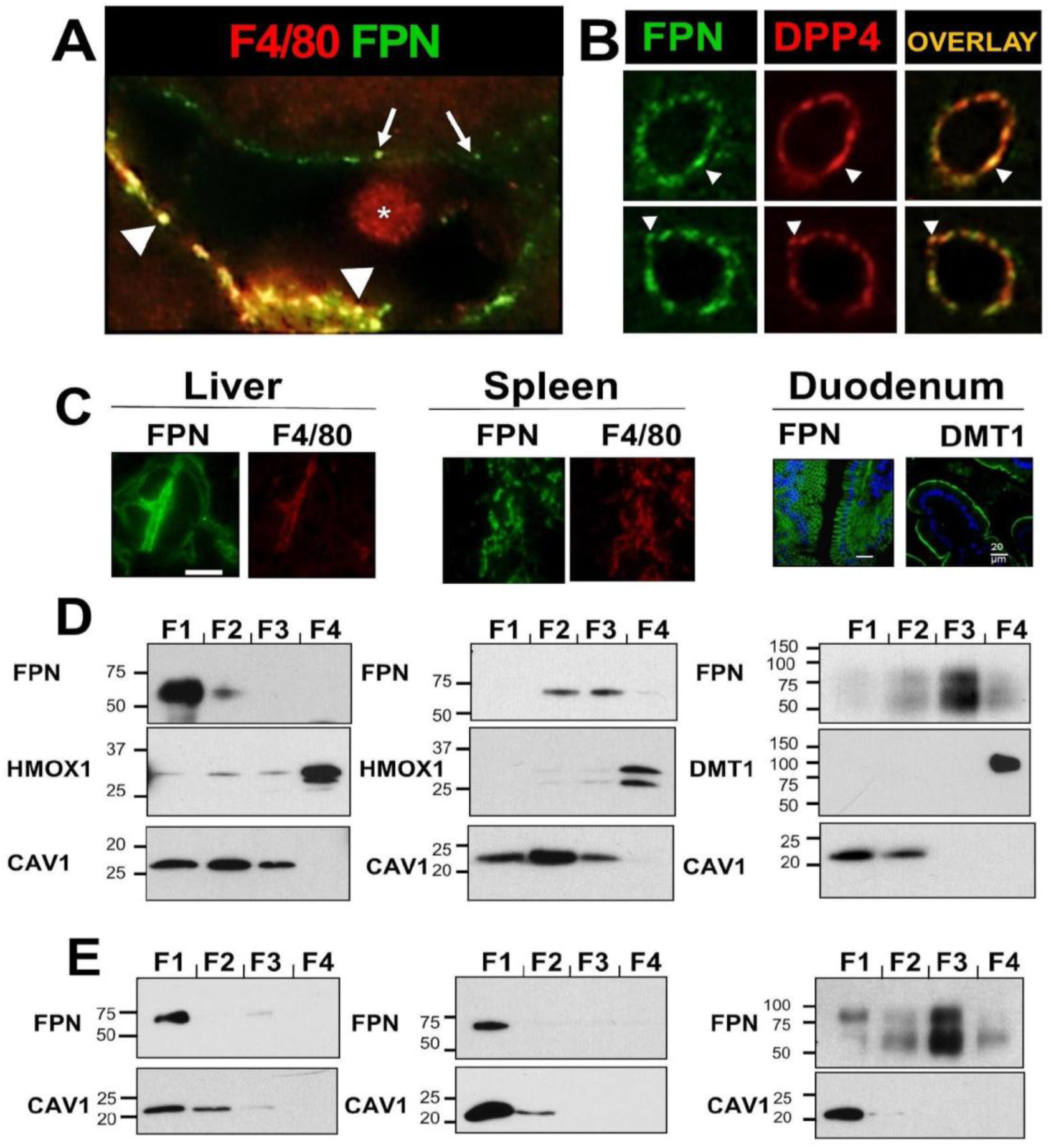
Enrichment of *Bmp6* KO hepatic ferroportin (Fpn) in detergent resistant membrane compartment. (A) Confocal analysis of Fpn staining with F4/80, indicating the enrichment of the iron exporter in membrane compartments at the cell surface of tissue macrophages (arrowheads) and hepatocytes (arrows) in *Bmp6* KO mice. **(B)** Confocal analysis of Fpn staining with Dipeptidyl peptidase-4 (Dpp4), showing the enrichment of Fpn in membrane compartments at the cell surface of hepatocytes (arrows) in *Bmp6* KO mice. **(C)** Immunohistofluorescence of Fpn in liver, spleen, and duodenum from *Bmp6* KO mice. F4/80 staining indicates Fpn expression in Kupffer cells and hepatocytes in the liver, and in splenic macrophages in the red pulp. In the duodenum, Fpn is strongly expressed at the basolateral membrane of enterocytes, opposite to the brush border localization of Dmt1. **(D & E)** Immunoblotting detection of Fpn, Hmox1, Dmt1, and caveolin1 (Cav1; raft marker) in detergent resistant membrane (DRM) and non-detergent resistant membrane (NDRM) fractions isolated from liver, spleen, and duodenum of 8 weeks-old *Bmp6* KO mice. Similar volumes (D) or similar amounts of protein (E) from pooled fractions A, B, C, and D were analyzed. Hepatic and splenic Fpn are enriched in fractions containing Cav1, while intestinal Fpn is not. The molecular weight marker’s position and size in kilodaltons (kDa) are indicated on the left.

## DISCUSSION

In this investigation, we examined the hepatic iron overload phenotype of *Bmp6* KO mice while concurrently exploring the expression pattern of the iron exporter Fpn at tissue, cellular, and subcellular levels. This approach has provided novel insights into the intricacies of liver iron homeostasis.

Initially, we identified some heterogeneity in the iron overload phenotype across distinct hepatic lobes. These findings underscore the necessity of ensuring uniformity in iron dosage or Perls staining during investigations of liver iron overload in *Bmp6* KO models. It is advisable to maintain consistency either by focusing solely on the same lobe or by utilizing homogenates of the entire liver for normalization purposes. Furthermore, we observed no discernible correlation between iron levels in each lobe at different ages and the mRNA expression levels of *Hamp* or *Fpn* which remains constant between liver lobes (not shown). The variations in iron accumulation between lobes in *Bmp6* KO mice can likely be attributed to a strong iron overload and anatomical considerations pertaining to the distribution of iron-enriched blood from the intestine. This blood enters the branches of the portal veins and hepatic artery, thereby likely contributing to the observed interlobe disparities.

Importantly, in 24-weeks aged *Hfe* KO mice (a model of hemochromatosis), no such difference in iron loading between lobes was observed ^14^. The varying levels of iron accumulation observed between Bmp*6* and *Hfe* KO mouse models may reflect differences in the severity or mechanisms of iron overload in response to genetic mutations. Indeed, the specificity of each iron overload model should be carefully considered when designing and interpreting experimental investigations.

Within *Bmp6* KO mice, the phenomenon of iron overload also displayed heterogeneity within the same hepatic lobe. In young mice and in specific areas of the liver section within each lobe (except for CL, harboring the lowest iron content), two distinct iron overload patterns emerged: areas of low iron overload (LI) exhibiting concentrated iron levels in the centrilobular zone, and high iron overload regions (HI) marked by discrete iron aggregates and lacking a defined zonation. Once more, this underscores the importance of exercising caution when conducting liver lobe analyses in this model and emphasizes the need to avoid sampling a portion of the lobe but rather examining the lobe in its entirety. As these mice aged, iron overload increased across all hepatic lobes, resulting in a loss of iron zonation. Critically, the liver iron zonation in LI corresponded to an opposing zonation of Fpn expression. Specifically, in LI, Fpn protein is predominantly detected in periportal hepatocytes and Kupffer cells, indicating distinct regulatory mechanisms governing Fpn expression in periportal versus centrilobular hepatocytes. Parallel observations were made in *Hamp* KO mice ^5^. This pattern of Fpn expression was similarly noted in rats at the mRNA level via *in situ* hybridization ^6^, suggesting contributions of transcriptional and post-transcriptional (mRNA stability) regulations to periportal zonal Fpn expression. However, the possibility of other regulatory mechanisms cannot be excluded. Remarkably, *Hamp* mRNA expression was localized within the same periportal zone where Fpn is expressed ^15^. Regulation of Hamp1 expression in response to body iron levels in the liver involves BMP ligand binding to their receptors, particularly BMP6, which triggers SMAD phosphorylation and gene expression regulation ^16^. Hemojuvelin (Hjv), a BMP co-receptor, enhances the SMAD phosphorylation pathway by binding to BMP ligands and receptors. Hamp expression is significantly reduced in the absence of Hjv ^17^. Notably, Hjv was found to be selectively expressed by periportal hepatocytes in mouse livers ^17^. Collectively, these findings suggest that the localized expression and function of Hamp in the periportal zone contribute to sustaining Fpn expression in this area (Fig.S5). Worth noting, our observations revealed that periportal zones contain a higher abundance of hepatic macrophages compared to centrilobular regions.

The study conducted by Latour et al. ^18^ presents a comprehensive analysis using a range of mouse models, including single and double knockouts of iron regulators. These models exhibit diverse iron overload patterns—periportal versus centrilobular—coupled with varying reductions in hepcidin expression. Upon examining the collective dataset, a clear correlation emerges between the detected level of hepcidin expression (mRNA) in the mouse models and the spatial distribution of Perls staining (indicative of iron levels) within the liver (Fig.S5). In instances where mice displayed extremely low hepcidin levels [Group D: *Bmp6* KO (male), *Hamp* KO, *Hjv* KO (male), *Bmp6* KO / *B2m* KO, *Bmp6* KO / *Tfr2* KO, and *Bmp6* KO / *B2m* KO / *Trf2* KO], iron deposition predominantly occurred in the centrilobular regions, aligning with the observations in this study (particularly evident in young *Bmp6* KO mice). In these mice, the notably diminished hepcidin levels exert dual positive impacts on both systemic (macrophages, duodenum) and local (hepatic) Fpn expression, consequently contributing to the redistribution of iron from periportal hepatocytes to hepatocytes encircling the central veins (Fig.6C and Fig.S5).

**Fig.6:**
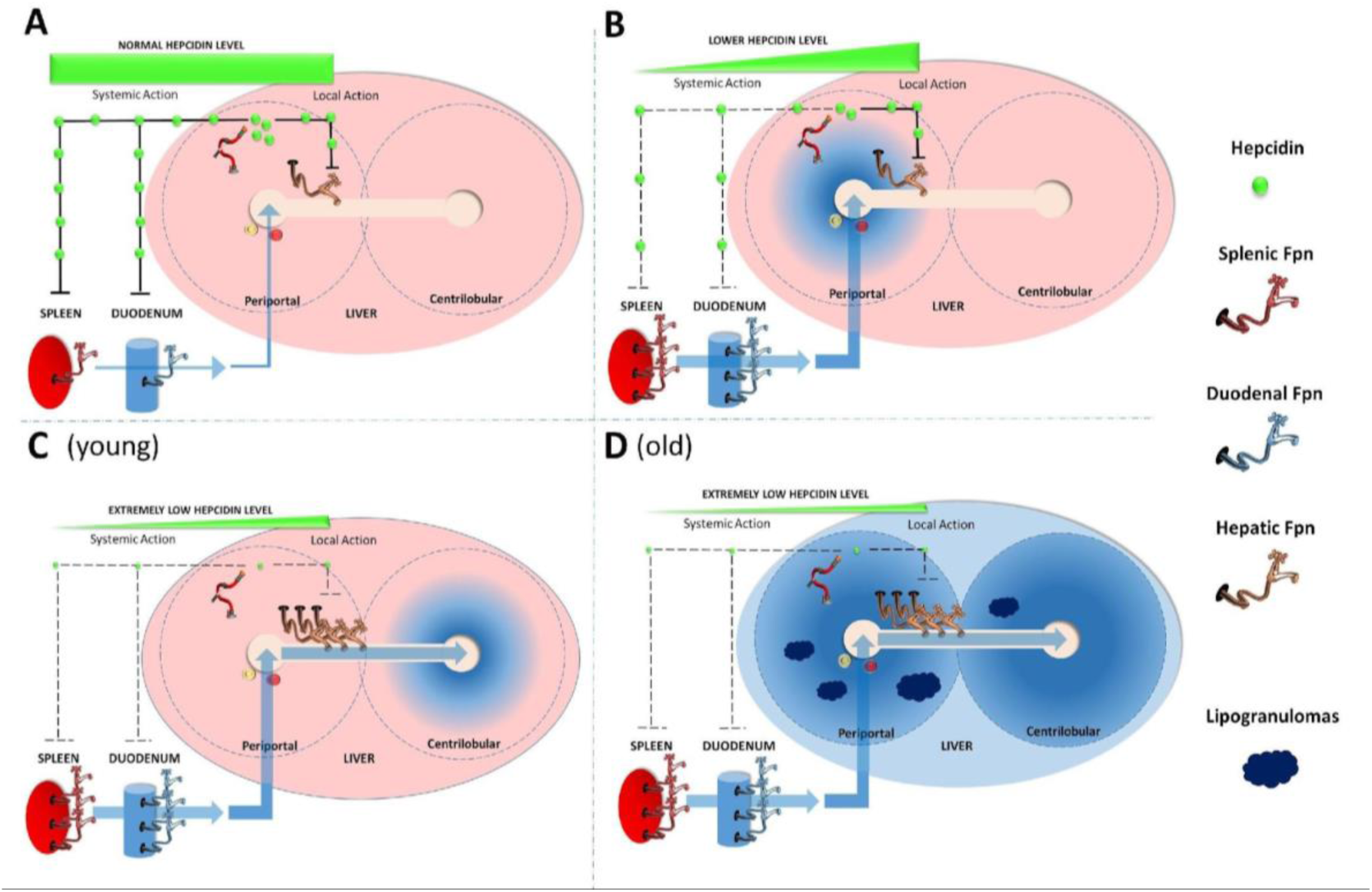
Influence of hepcidin and ferroportin centrilobular expression on hepatic iron distribution. (A) In most wildtype mice like C57BL6 with normal hepcidin levels, both local (hepatic) and systemic (splenic and intestinal) effects of Hamp prevent liver iron overload. (B) A modest decrease in Hamp levels, observed in CD1 wildtype mice, *Hfe*, *B2m*, and *Tfr2* knockout mice, heterozygote Hamp +/- mice, *Bmp6* and Hjv knockout females, and Neogenin mutants, leads to iron accumulation primarily in the periportal area. Reduced systemic Hamp action results in increased iron recycling by macrophages and intestinal absorption, leading to iron accumulation in periportal hepatocytes due to the maintained downregulation of hepatic Fpn. **(C)** A dramatic decrease in Hamp levels impairs both systemic and local Hamp actions, leading to strong Fpn expression in tissue macrophages, enterocytes, and periportal hepatocytes. This results in increased iron recycling by macrophages and intestinal absorption, leading to iron flux through the periportal vein. Due to high Fpn expression in periportal hepatocytes, iron is mainly redistributed to the centrilobular area, where hepatocytes accumulate iron. This iron zonation is observed in *Bmp6* and Hjv knockout males, Hamp knockout mice, and double or triple knockouts (*Bmp6*/B2m, *Bmp6*/Tfr2, *Bmp6*/B2m/TfR2). (D) In contexts of extremely low Hamp expression, iron zonation tends to disappear over time, leading to the formation of lipogranulomas predominantly in the periportal zone.

Conversely, in *Bmp6* KO females (Fig.S5; Group C), who exhibit slightly elevated hepcidin levels compared to *Bmp6* KO males, iron distribution predominantly occurs within the periportal regions (Fig 6B) (18). Similarly, zonal patterns of injury, characterized by periportal hemosiderin deposition, are observed in *Hfe* KO mice and individuals with hemochromatosis ^19,20^. Despite expressing hepcidin, albeit at inappropriately low levels relative to their body iron stores ^21–24^, both *Hfe* KO mice and hemochromatosis patients experience such periportal iron overload. This phenomenon extends to mice with low or inadequate hepatic Hamp expression, encompassing various genotypes [*Bmp6* KO (female), *Hfe* KO, *B2m* KO, *TFR2* KO, *Hamp* +/-, *Hjv* KO (females), *Neogenin* KO], and promote intestinal iron absorption and heme iron recycling from the spleen while preserving a local regulation of hepatic Fpn (Fig.6B; Fig.S5). In such scenarios, a continuous influx of iron-rich blood from the intestines via the portal vein, coupled with minimal fluctuations in periportal Fpn repression, facilitates iron accumulation within hepatocytes surrounding the periportal veins. Moreover, we demonstrate that CD1 WT mice exhibit modest iron accumulation in periportal hepatocytes, a phenomenon not observed in other wildtype strains like C57BL/6 (no discernible iron accumulation, data not shown). Once again, CD1 mice were shown to express less Hamp than other WT mice (personal communication). Consequently, contingent on the threshold of hepcidin levels, iron accumulation occurs either within the periportal or centrolobular zones (Fig 6 & Fig.S5). As time progresses, this distinctive zonal distribution diminishes, resulting in a more diffuse distribution of iron across the entire liver (Fig.6D).

The liver, an intricate organ, encompasses a diverse array of metabolic pathways. To ensure its optimal functionality, metabolic zonation is imperative ^25^, and for this purpose, the liver parenchyma exhibits notable heterogeneity and functional adaptability. This demand for spatial organization appears to extend to hepatic iron metabolism. The redistribution of iron from periportal to centrilobular zones, facilitated by the gradient of Fpn expression, could constitute a protective mechanism aimed at limiting hepatic injury. Indeed, the centrilobular area is acknowledged as a detoxification zone characterized by reduced oxygenation, potentially rendering centrilobular hepatocytes less prone to iron and hemosiderin accumulation. Pathological iron accumulation within tissues exacerbates the generation of reactive oxygen species (ROS) and triggers detrimental effects primarily associated with oxidative stress. Compared to centrilobular cells, periportal hepatocytes contain a substantial number of peroxisomes and mitochondria, significant sources of reactive oxygen species (ROS) ^25^. In order to mitigate the impact of ROS, these cells also exhibit elevated levels of superoxide dismutase 2, an enzyme responsible for generating H_2_O_2_, a potent catalyst for dangerous free radicals formed in conjunction with ferrous iron through the Fenton reaction. Consequently, the accumulation of iron within cells situated in regions characterized by lower oxygenation, such as the centrilobular area, might potentially confer reduced harm to the overall organism.

Interestingly, as time progressed, the prominence of hepatic lesions, characterized as lipogranulomas, exhibited a more marked manifestation in periportal zones in *Bmp6* KO mice. These lesions were discerned as substantial clusters of macrophages displaying elevated levels of Fpn and containing significant quantities of both hemosiderin and ceroid/lipofuscin. Ceroid, akin to lipofuscin, constitutes a yellow-brown pigment originating from an insoluble polymer of oxidized lipids and proteins. It is recognized as a pathological pigment formed through the phagocytosis of lipids from deceased hepatocytes (exogenous lipids/heterophagy), subsequently accumulating in hypertrophied macrophages following liver injury. Interestingly, in old mice exhibiting increased erythrophagocytosis and iron overload in the spleen, similar large iron-rich aggregates were observed through tissue staining inside some morphologically damaged macrophages ^26^. In addition to these intracellular aggregates, some large amorphous extracellular aggregates were also detected, likely emerging from damaged red pulp macrophages.

As previously discussed, elevated iron deposition in periportal hepatocytes could trigger a potent oxidative stress reaction culminating in cell death. In certain murine models of hemochromatosis, ferroptosis — a specific form of cell death instigated by iron-dependent lipid peroxidation — has been observed in hepatocytes ^27^. Intriguingly, periportal macrophages have been found to be more prone to phagocytosis in comparison to their centrilobular counterparts^28^. In addition, we observed the presence of more macrophages in periportal zone than in the centrolobular one. Moreover, it is plausible that the composition of lipogranulomas in *Bmp6* KO involves distinct macrophage populations. In healthy liver, Kupffer cells do not solely constitute the recognized tissue macrophages, and during liver diseases, additional macrophage subsets are recruited, such as the lipid-associated macrophages (LAM) ^29^. Consequently, it can be inferred that lipogranulomas in *Bmp6* KO likely encompass resident and/or recruited macrophages laden with iron and lipids, acquired through the phagocytosis of sideronecrotic and ferroptotic hepatocytes.

The iron exporter Fpn was found to be strongly present at the membranes of Kupffer cells and on the surface of hepatocytes. However, Fpn was not detected on the membrane that lines the biliary duct, suggesting that the transporter does not directly transport iron for biliary secretion. Despite earlier evidence of Fpn mRNA expression in hepatic stellate cells (HSC) ^6^, we did not observe Fpn protein expression in these cells.

At the subcellular level, in both Kupffer cells and hepatocytes, the iron exporter was present in discrete domains in the plasma membrane. In hepatocytes, Fpn was shown to strongly colocalize with Dpp4 (dipeptidyl peptidase-4 aka CD26), a cell surface glycoprotein. Interestingly, Dpp4 interacts with Caveolin-1 which facilitates its recruitment into lipids rafts in T cells ^12^. Furthermore, in the hematopoietic environment, Dpp4 activity is primarily located with lipid rafts ^13^. Lipid rafts correspond to specific molecular platforms, floating at the cell surface and involved in numerous biological functions such as cell signaling ^30–32^. In cultured macrophages, Fpn has been found to be highly concentrated within lipid rafts, a crucial localization for the regulation of the iron exporter by Hamp ^11^. For the first time, our study demonstrates that both hepatic cells (hepatocytes and Kupffer cells) as well as splenic macrophages exhibit Fpn enrichment within lipid rafts in an *in vivo* context. These findings suggest that Fpn resides within specific subdomains of the plasma membrane in both hepatocytes and tissue macrophages (Kupffer cells and splenic macrophages). In contrast, the form of Fpn found in the duodenum, expressed at the basolateral membrane of enterocytes, is predominantly detected in a non-raft fraction. These observations emphasize the existence of differences between duodenal, hepatic, and macrophage Fpn, a phenomenon previously noted in relation to post-transcriptional modifications ^8^. It is plausible to speculate that these observed differences, including lipid raft localization and glycosylation, could play a significant role in the iron exporter’s transport activity and/or its regulation. Notably, the degradation of Fpn induced by Hamp appears to be more efficient in macrophages compared to enterocytes, which are more resilient to Hamp’s effects ^33–35^. Intriguingly, during the suckling period, enterocyte ferroportin seems less responsive to the degradative impact of circulating hepcidin ^36^. This finding correlates with a change in the electrophoretic mobility of Fpn between suckling and adult mice. In suckling mice, the ferroportin protein is smaller, which could be indicative of an alternatively spliced form of the transporter or different post-translational modifications, such as glycosylation. Subsequent investigations are necessary to elucidate the biological implications underlying the disparities observed among the various cellular forms of Fpn.

## CONCLUSION

In summary, our exploration of hepatic iron regulation in *Bmp6* KO has unveiled intricate patterns of ferroportin (Fpn) expression and its impact on zonal iron distribution in the liver. It reveals the detailed aspects of Fpn localization, from its specific presence in Kupffer cells and hepatocyte apical membranes within periportal zones to its association with macrophages in iron-rich regions and hepatic lesions. Finally, our study underscores the importance of understanding Fpn’s subcellular localization for comprehending its transport activity and its modulation by hepcidin.

## Supporting information

Supplemental Data BMP6KO

## AUTHORSHIP

### Contribution

C.B., A.W., and L.R. designed protocols, conducted experiments, and analyzed data (CB: Perls staining, RT-PCR & immunofluorescence, A.W: western blot and immunofluorescence; L.R.: lipid raft studies). C.L. carried out experiments (mice genotyping and RT-PCR) and analyzed data. H.C. and M-P.R. participated in data discussions and manuscript review. F.C-H. formulated protocols, conducted experiments (western blot and IF), critically evaluated the data, and wrote the paper. The correction and improvement of the English language in this article were assisted by ChatGPT, a language model developed by OpenAI.

## ACKNOWLEDGEMENTS

We wish to acknowledge Ophélie Gourbeyre and Benjamin BILLORE for technical assistance.

