## Supplemental Data BMP6KO for "NEW INSIGHTS INTO THE HEPATIC IRON PHENOTYPE OF BMP6 KNOCKOUT MICE"



### SUPPLEMENTAL DATA & LEGENDS

#### Table S1 - Sequence of primers used in quantitative PCR.

| Gene | Sequence | Primer sense |
|---|---|---|
| *Hprt* | CTGGTTAAGCAGTACAGCCCCAA | Forward |
| | CAGGAGGTCCTTTTCACCAGC | reverse |
| *Fpn (Scl40a1)* | CATTGCTGCTAGAATCGGTCTT | forward |
| | GCAATCGTGTCACCGTCAAAT | reverse |
| *Hamp* | AAGCAGGGCAGACATTGCGAT | forward |
| | CAGGATGTGGCTCTAGGCTATGT | reverse |

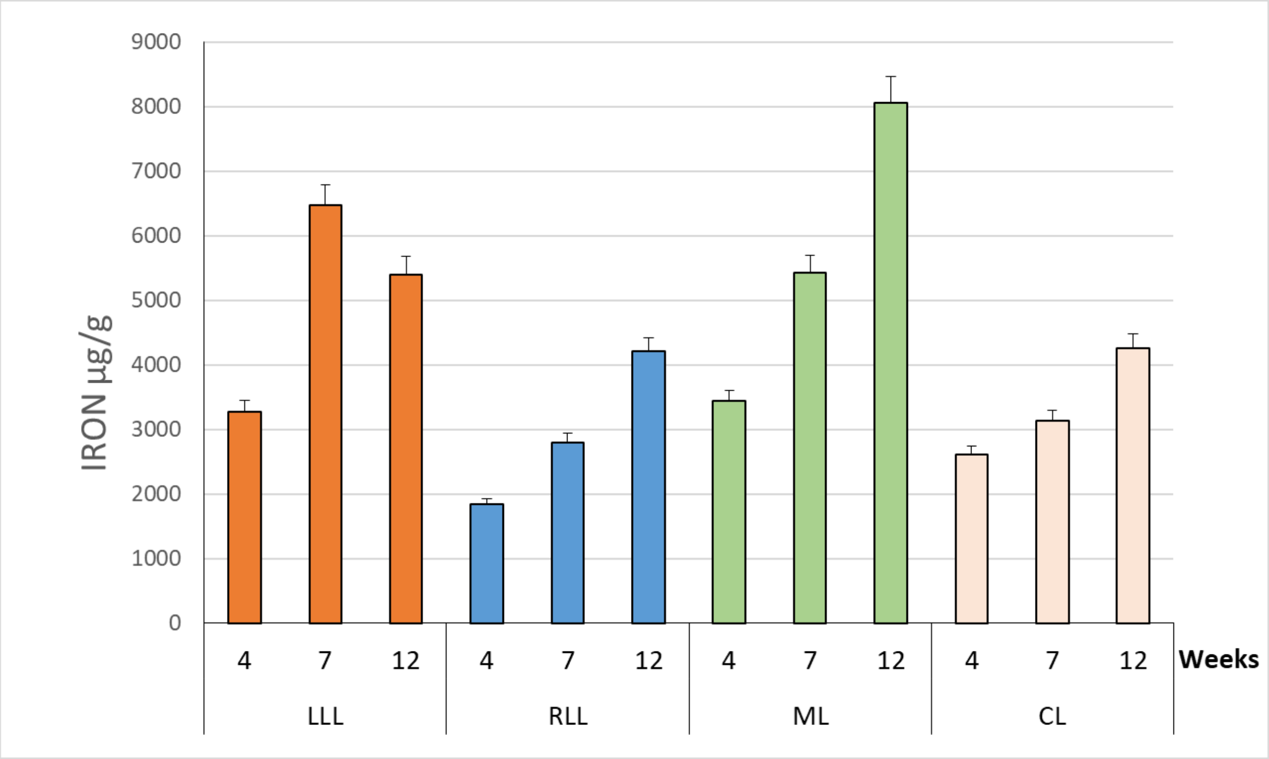


**Fig.S1**: **Quantitative assessments of nonheme iron in hepatic lobes** by pooling distinct lobes [left lateral lobes (LLL), right lateral lobes (RLL), medium lobes (ML), and caudate lobes (CL)] from three mice per genotype.  Iron measurements were conducted at various ages (4, 7 and 12 weeks) with the iron content of pooled lobes measured three times (three technical replicates). ML and, to a lesser extent, LLL were found to accumulate more iron than the RLL and CL. These measurements support the observations from the Perl's staining (Figure 1). Collectively, our findings indicate differences in iron accumulation among lobes in Bmp6 KO mice.

Céline BESSON[1], Alexandra WILLEMETZ[2,3], Chloé LATOUR[1], Lorenne ROBERT[4], Helene COPPIN[1], Marie-Paule ROTH[1] and François CANONNE-HERGAUX[1, #]


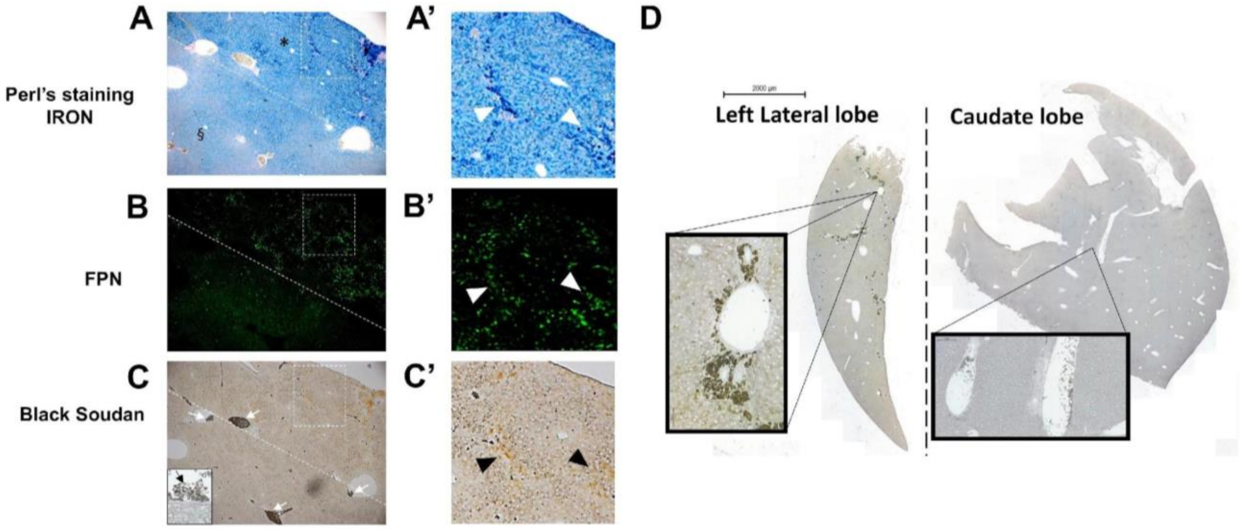


**Fig.S2: Perls Prussian blue (A & A'), ferroportin (Fpn, B & B'), and Soudan black (C & C') staining of hepatic left lateral lobe (LLL) from 8-week-old *Bmp6* KO mice**. Iron localization was heterogeneous, with some zones showing lower (§) and others showing higher iron accumulation (*). Iron accumulation was associated with strong Fpn staining (arrowheads in B') and the presence of ceroid/lipofuscin accumulation (arrowheads in C'). Arrows in C and inset in C showed dark gray staining of red blood cells with Soudan black in hepatic vessels. (D) Soudan black staining of the left lateral (LLL) and caudate (CL) lobes of an 8-week-old *Bmp6* KO mouse. Only the LLL presents ceroid/lipofuscin accumulation.

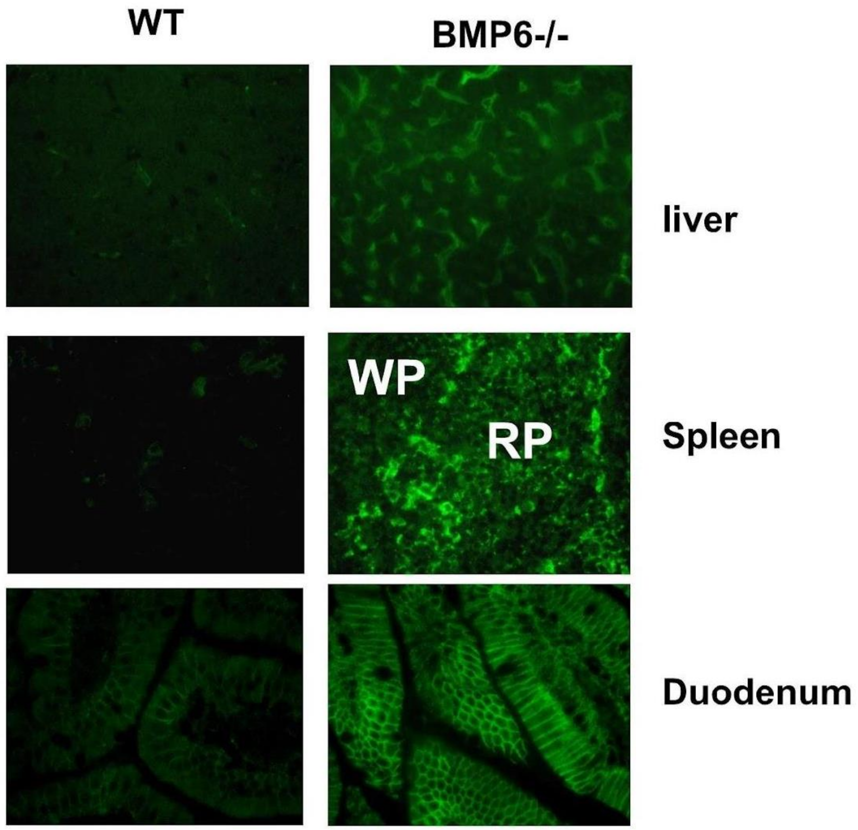


**Fig.S3: Immunohistofluorescence of Fpn in wildtype (WT) and *Bmp6* knockout (KO) mice.** **E**ight-week-old WT and *Bmp6* KO mice were sacrificed, and tissues (liver, spleen and duodenum) were processed for the detection of Fpn by immunofluorescence (green). Fpn expression was markedly increased in *Bmp6* KO mice compared to WT.

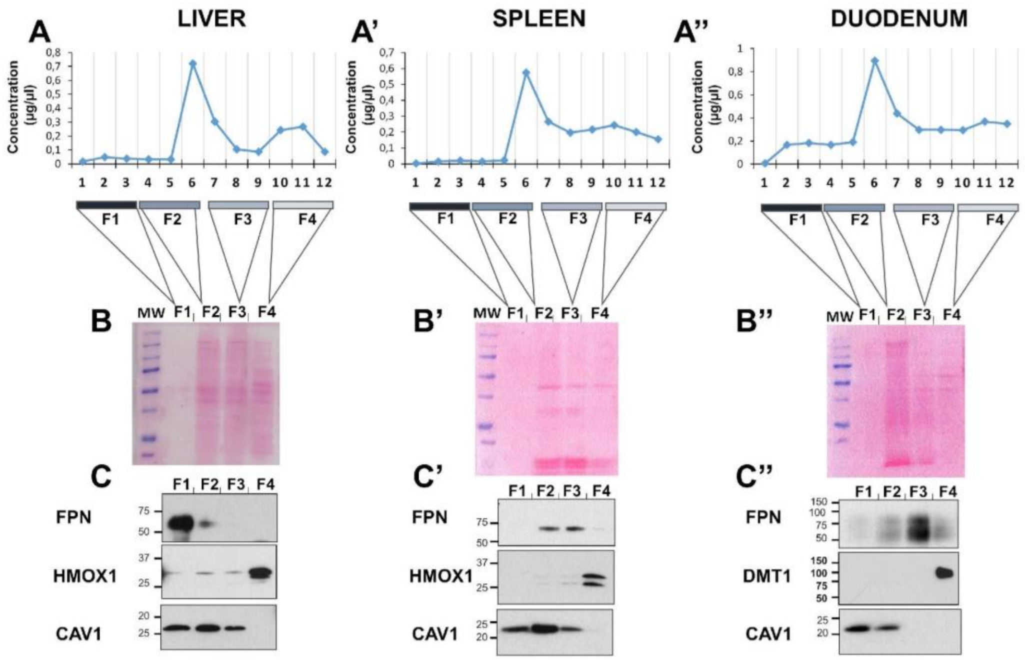


**Fig.S4: Analysis of the iodixanol gradient fractions isolated from *Bmp6* KO tissues.** (A, A', and A") Protein quantification of each fraction collected from the iodixanol gradient after ultracentrifugation of liver (A), spleen (A'), and duodenum (A") membrane protein extracts. Samples were combined in groups of three for F1 (fractions 1, 2, and 3), F2 (fractions 4, 5, and 6), F3 (fractions 7, 8, and 9), and F4 (fractions 10, 11, and 12) before gel electrophoresis and western blot analysis. (B, B', and B") Ponceau S staining of the PVDF membrane showing the protein distribution after transferring F1, F2, F3, and F4 extracts. (C, C', and C") Immunoblotting detection, as presented in Fig. 5B, of Fpn, Hmox1, Dmt1, and Cav1 in F1, F2, F3, and F4 fractions isolated from liver (C), spleen (C'), and duodenum (C"). The position and size in kilodaltons (kDa) of the molecular weight markers (MW) are indicated on the left. Notably, hepatic Fpn is significantly enriched with Cav1 in F1, which is the pooled fraction with the lowest protein content.

Céline BESSON[1], Alexandra WILLEMETZ[2,3], Chloé LATOUR[1], Lorenne ROBERT[4], Helene COPPIN[1], Marie-Paule ROTH[1] and François CANONNE-HERGAUX[1, #]


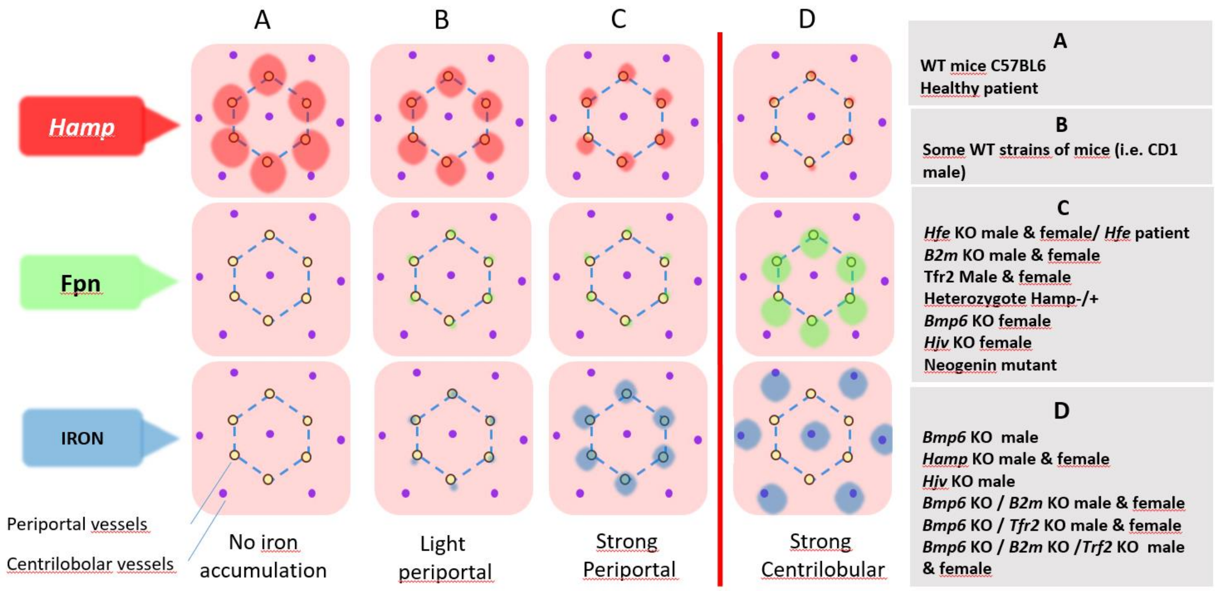


**Fig.S5: Schematic illustration depicting the expression and localization of iron, ferroportin, and hepcidin in various mouse models of iron overload.** This figure illustrates the varying levels of hepcidin and Fpn in different mouse models (A to D), leading to varying degrees and localization of liver iron. A crucial threshold (depicted by the red line) is observed, beyond which the iron distribution transitions from periportal to centrolobular areas.
